# Direct anabolic three carbon metabolism via propionate to a six carbon metabolite occurs in human heart and in vivo across mouse tissues

**DOI:** 10.1101/758706

**Authors:** Mary T. Doan, Michael D. Neinast, Erika L Varner, Kenneth Bedi, David Bartee, Helen Jiang, Sophie Trefely, Peining Xu, Jay P Singh, Cholsoon Jang, Eduardo Rame, Donita Brady, Jordan L. Meier, Kenneth Marguiles, Zoltan Arany, Nathaniel W. Snyder

## Abstract

Anabolic metabolism of carbon in mammals is mediated via the one and two carbon carriers S-adenosyl methionine and acetyl-coenzyme A (acetyl-CoA). In contrast, anabolic metabolism using three carbon units via propionate is not thought to occur. Mammals are primarily thought to oxidize the 3-carbon short chain fatty acid propionate by shunting propionyl-CoA to succinyl-CoA for entry into the TCA cycle. We found that this may not be absolute and that in mammals one non-oxidative fate of two units of propionyl-CoA is to condense to a six-carbon trans-2-methyl-2-pentenoyl-CoA (2M2PE-CoA). We confirmed this pathway using purified protein extracts provided limited substrates and confirmed the product with a synthetic standard. In whole-body *in vivo* stable isotope tracing with infusion of ^13^C-labeled valine achieving steady state, 2M2PE-CoA formed via propionyl-CoA in multiple murine tissues including heart, kidney, and to a lesser degree in brown adipose tissue, liver, and tibialis anterior muscle. Using *ex vivo* isotope tracing, we found that 2M2PE-CoA formed in human myocardial tissue incubated with propionate to a limited extent. While the complete enzymology of this pathway remains to be elucidated, these results confirm the *in vivo* existence of at least one anabolic three to six carbon reaction conserved in humans and mice that utilizes three carbons via propionate.

**Highlights:**

- Synthesis and confirmation of structure 2-methyl-2-pentenoyl-CoA
- ***In vivo*** fate of valine across organs includes formation of a 6-carbon metabolite from propionyl-CoA
- ***Ex vivo*** metabolism of propionate in the human heart includes direct anabolism to a 6-carbon product
- In both cases, this reaction occurred at physiologically relevant concentrations of propionate and valine
- ***In vitro*** this pathway requires propionyl-CoA and NADH/NADPH as substrates

## Introduction

One-and two-carbon (1C and 2C) metabolism are highly conserved metabolic processes for anabolic and catabolic biotransformation. 1C metabolism is facilitated via folate intermediates, with a formyl-group transferred through activated intermediates including S-adenosyl methionine (1). 2C metabolism occurs primarily via the formation of the thioester acetyl-coenzyme A (acetyl-CoA) and transfer of the activated acetyl-group from the thioester to substrates (2, 3). Both 1C and 2C metabolic pathways are conserved across bacteria, yeast, plants, and animals. In contrast, direct anabolic three-carbon (3C) metabolism is thought to occur in a more limited set of organisms, including in propionic bacteria (4), and in metabolism of branched chain hydrocarbons and hormones in some insects (5). It is generally thought that mammals do not use 3C intermediates directly for anabolic reactions, with the notable exception of pyruvate as a gluconeogenic substrate (6). In propionate mediated 3C metabolism, propionyl-CoA, the CoA thioester of propionic acid, functions as the central metabolic intermediate. For instance, propionyl-CoA is an intermediate in the catabolism of two branched amino acids (BCAAs), isoleucine and valine (7). Propionyl-CoA can also be formed from catabolism of diverse substrates including odd-chain fatty acids, cholesterol, C-5 ketone bodies, threonine, and methionine (5). Since it is energetically unfavorable to metabolize propionyl-CoA via β-oxidation, the canonical use of propionyl-CoA is for anaplerosis into the TCA cycle by metabolism to succinyl-CoA via D-then L-methylmalonyl-CoA (8, 9).

Propionate metabolism in humans results in unique biochemical effects with mostly unknown mechanisms of action. Infusion or feeding of propionate causes perturbations of liver metabolism in rodents, dogs, and humans (10, 11). Extreme pathological defects in propionate metabolism including propionic acidemia (PA), an inherited disorder caused by an aberrant function of propionyl-CoA carboxylase (PCC), result in significant morbidity and mortality (12) and systemic metabolic dysregulation (13). Although PA is often detectable by newborn screening, even after diagnosis, prognosis remains poor due to sub-clinical metabolic dysregulation, life-threatening metabolic decompensations, and sequelae from severe crisis (14). Especially during these metabolic crisis, patients with PA have high levels of organic acids and acylcarnitines in the blood and urine which include not only propionic acid/propionylcarnitine but also a wider dysregulated metabolome of mostly unidentified metabolites (15). This suggests that propionate can be metabolized to a wide number of poorly described products *in vivo*. The relevance of these metabolites to normal physiology with lower levels of circulating propionate is unknown. Furthermore, since anaplerosis to the TCA cycle is tightly regulated, the non-oxidative fates of propionyl-CoA that is not shunted through succinyl-CoA is unclear.

Recently, we described a potential route for anabolic 3C metabolism in cell culture and *ex vivo* platelets to convert propionyl-CoA into the acyl-CoA thioester of *trans*-2-methyl-2-pentenoic acid (2M2PE), a 6-carbon compound that retained all of the carbons from two monomeric units of propionate (16). This contrasts with the monomethyl branched chained fatty acids (mmBFCAs) synthesis in mammalian adipose tissues, which requires 2C metabolism via acetyl-CoA (17). However, the existence of this pathway in intact tissues and more physiologically relevant systems remained unknown. Here, we confirmed this pathway by using purified protein extracts and a limited set of substrates, and we compared the product against a synthetically prepared standard. We then tested the existence of this pathway *in vivo* in mice using an infusion of U-^13^C-valine which maintains 3 labeled atoms in catabolism to propionyl-CoA (**Fig. 1**) (18). Steady state infusion of valine in awake and mobile rodents was chosen for physiological relevance to avoid induction of artificial metabolism from supraphysiological propionate concentrations while maintaining normal oxidative uses of propionyl-CoA. Finally, we examined the formation of 2M2PE-CoA from ^13^ C_3_-propionate *ex vivo* using a mismatched human donor heart.

**Figure 1.**
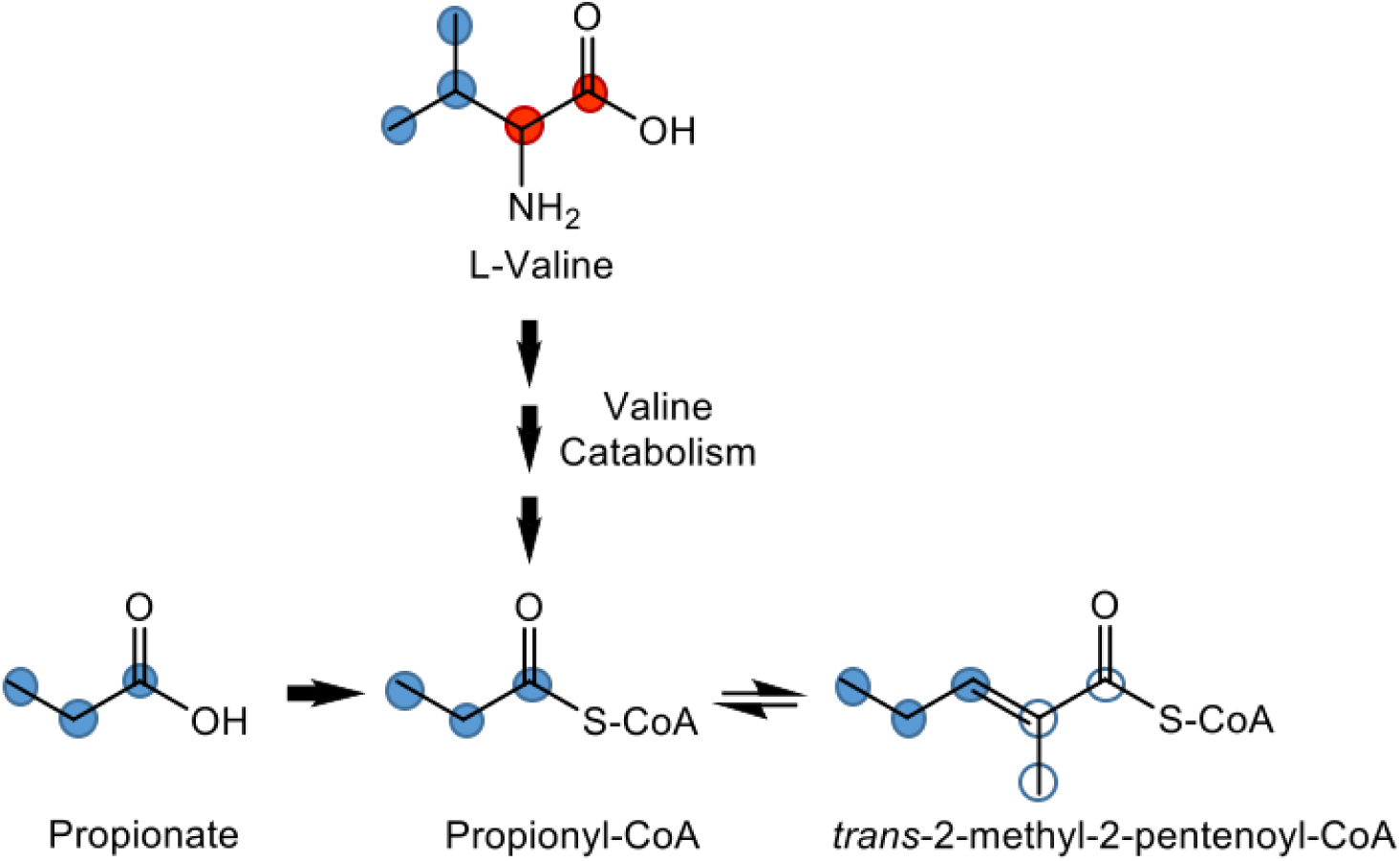
Isotope incorporation into propionyl-CoA and 2M2PE-CoA from U-^13^C-valine maintains integers of 3 contiguous carbon atoms. Carbon-13 from U-^13^C-valine or U-^13^C-propionate is indicated by red, blue, and blue open circles. Catabolism of valine or activation of propionate produces [^13^C_3_]-propionyl-CoA which retains 3 labeled atoms (blue). Condensation of 2 molecules of propionate results in the formation of *trans*-2-methyl-pentenoyl-CoA with [^13^C_3_] labeling of all six-carbon atoms: 3 deriving from one propionyl-CoA (blue) and 3 deriving from a second propionyl-CoA (blue open).

## Methods

### Materials and reagents

[^13^C_3_]-sodium propionate was purchased from Cambridge Isotopes (Tewksbury, MA). 5-sulfosalicylic acid (SSA), trichloroacetic acid, and ammonium acetate were purchased from Sigma-Aldrich (St. Louis, MO). Optima LC-MS grade methanol (MeOH), acetonitrile (ACN), formic acid, and water were purchased from Fisher Scientific (Pittsburgh, PA).

### Synthesis of 2-methyl-2-pentenoyl-CoA

Synthesis was conducted as previously described with modifications (19). Briefly, diisopropylethylamine (0.12 mL, 0.70 mmol) was added to a solution of 2-methyl-2-pentenoic acid (30.6 mL, 0.26 mmol) in dichloromethane (2 mL) cooled to 0 °C. Isobutyl chloroformate (64 uL, 0.65 mmol) was added at 0 °C with stirring. The resulting solution was allowed to warm to 22 °C and stirred for an additional 2 h, after which, volatiles were removed under reduced pressure to provide a colorless residue. Tetrahydrofuran (THF, 1 mL) was added to the residue resulting in precipitation of diisopropylethylammonium chloride. This mixture was transferred to a 1.6 mL conical tube and centrifuged to pellet the solid. The THF solution was added to a solution of coenzyme A (30 mg, 0.039 mmol) in aqueous bicarbonate solution (1 mL, 500 mM, pH 8). The biphasic mixture was stirred vigorously at 22 °C. After 3 h, stirring was stopped and THF was evaporated under a stream of N_2_. The remaining aqueous solution was neutralized with trifluoroacetic acid (TFA, 20 mL) and immediately purified by C_18_-reversed phase-silica chromatography. Method: (solvents - A: 0.05% TFA in water, B: acetonitrile) gradient - 0-2 column volumes (CV), 0% B; 2-12 CV, 0-50% B; 12-13 CV, 50-100% B; and 13-25 CV, 100% B. Fractions containing the desired product were pooled and lyophilized to provide a fluffy white powder (17.7 mg, 52.5% yield). ^1^H NMR (500 MHz, D_2_O) δ 8.63 (s, 1H), 8.42 (d, *J* = 3.2 Hz, 1H), 6.77 (t, *J* = 7.2 Hz, 1H), 6.18 (d, *J* = 5.7 Hz, 1H), 4.91 (s, 1H), 4.89 – 4.84 (m, 1H), 4.60 (s, 1H), 4.35 – 4.19 (m, 2H), 4.01 (s, 1H), 3.89 (d, *J* = 9.1 Hz, 1H), 3.65 (d, *J* = 9.2 Hz, 1H), 3.44 (t, *J* = 6.8 Hz, 2H), 3.33 (t, *J* = 6.5 Hz, 2H), 2.99 (t, *J* = 6.3 Hz, 2H), 2.43 (q, *J* = 6.3 Hz, 2H), 2.19 (p, *J* = 7.5 Hz, 2H), 1.77 (s, 2H), 0.99 (t, *J* = 7.6 Hz, 3H), 0.94 (s, 3H), 0.82 (s, 3H). HRMS (ESI) calculated for C_27_H_44_N_7_O_17_P_3_S [M + H]^+^ 864.1800, found 864.1811.

### Protein purification and enzyme assay

Cells were passaged every 2-3 days at 80% confluence. Human hepatocellular carcinoma (HepG2) cells were cultured in high glucose DMEM (Thermo-Fisher Scientific) and supplemented with 10% fetal bovine serum (Gemini Biosciences) and penicillin/streptomycin (Thermo Fisher Scientific). SILEC labeling of HepG2 cells was achieved as previously described (20). As positive controls, 2M2PE-CoA was generated in cell culture, as previously described (16), by treating HepG2 cells for 1 hour with either 100 µM propionoate, ^13^C_1_-propionoate, or trans-2-methyl-2-pentenoic acid. Following the incubation, the acyl-CoA thioesters were extracted and analyzed.

To obtain purified protein, approximately 3 × 10^6^ HepG2 cells were collected and lysed by shaking in 500 µL of ice-cold FPLC buffer (50mM Tris/HCl pH: 7.5, 150mM NaCl, 100mM KCl, 1% NP-40 and 1% Glycerol) for 5 minutes. The cell debris was pelleted by centrifugation at 17,000 g, 4°C for 5 minutes. The 3 mg/mL protein sample was injected onto NGC Liquid Chromotography System (Bio-Rad) equipped with a HiPrep Sephacryl S-500 HR column equilibrated in FPLC buffer and separated into 156, 1 mL fractions. A 200 µL aliquot of each protein fraction was incubated at 37°C for 2 hours with 100 µL of the FPLC buffer containing propionyl-CoA (100 µM), NADH (1 mM), and NADPH (1 mM). Absorbance was recorded at 260 nm and 340 nm before and after the 2-hour incubation. Following the incubation, the acyl-CoA thioesters were extracted and analyzed.

### Tissue studies

As described previously (18), ^13^C-valine infusion was performed on 12-week old male C57/blk6 or db/db mice with a catheter surgically implanted on the right jugular vein. The mouse infusion setup included a tether and swivel system which permitted the mice to have free movement in the cage with bedding materials and free access to water. Briefly, the mice fasted for 5 hours and then infused intravenous (I.V.) with [U^13^C]-valine at ∼20% of the endogenous rate of appearance for 120 min to achieve steady state labeling. Unlabeled leucine and isoleucine were also infused with the labeled valine. Mice were sacrificed by pentobarbital injection followed by immediate cervical dislocation, and tissues were quickly harvested and snap-frozen within 1.5 min of cervical dislocation. Frozen brown adipose tissue (BAT), heart, kidney, liver, tibialis anterior muscle (TA), and gonadal white adipose tissue (gWAT) were individually cut over dry ice. Tissue samples were weighed to aliquots between 7 to 16 mg in 1.5 ml Eppendorf tubes then taken for extraction and analysis in conditions identical to those used for heart tissues. All animal work was approved by the University of Pennsylvania Institutional Animal Care and Use Committee.

Procurement of human myocardial tissues came from a healthy male donor on no medications and had no positive serologies. The decedent was 52 years old, weighed 90 kg, was 175 cm tall with a body mass index of 29.3 and body surface area of 20.9. Patient’s heart (352 g, left ventricle mass 213) displayed normal values of echocardiographic measurements. There were also no troponin leaks suggesting no ischemia. However, patient was turned down from transplantation due to a 50-60% stenotic left anterior descending coronary artery (seg 6) and died from a cerebrovascular accident/stroke of the middle cerebral artery. Otherwise, the donated heart was considered healthy enough to reflect normal human physiology. Heart tissues from a section of the left ventricle wall were individually cut over dry ice and weighed to aliquots between 40 to 55 mg in 1.5 ml Eppendorf tubes. For time course studies, prewarmed 1 ml of 100 µM [^13^C_3_]-sodium propionate in Tyrode’s buffer with 5mM glucose was added to each sample then incubated at 37°C for 15, 30, and 60 min. Samples were centrifuged at 10,000 x *g* for 2 min at 4°C to pellet the tissues. The resulting pellet was extracted and analyzed.

### Extraction of acyl-CoAs

Extractions for short chain acyl-CoAs were performed as previously described (16). Briefly, 1 mL of ice-cold 10% (w/v) trichloroacetic acid was added to tissues pre-weighed in a 1.5 mL Eppendorf tube, 50 µL of mixed acyl-CoA internal standard generated as previously described (20) was added, then the sample was vortexed 5 sec to mix. With a probe tip sonicator (Fisher), samples were homogenized and dismembranated by 12 × 0.5 sec pulses and centrifuged at 15,000 x *g* for 10 min at 4°C. The supernatant was purified by a solid-phase extraction on Oasis HLB 1cc (30 mg) solid phase extraction columns (Waters). Columns were pre-conditioned with 1 mL MeOH then 1 mL water, 1 mL of sample was loaded, the column was washed with 1 mL water, and then the elutate was collected into glass tubes using 1 mL of 25 mM ammonium acetate in MeOH. Eluates were evaporated to dryness under nitrogen gas and resuspended in 50 μl of 5% (w/v) SSA. Injections of 5 μL were made for LC-HRMS analysis.

### LC-HRMS and LC-MS/HRMS analysis

Sample analysis was performed as previously described (21). Briefly, acyl-CoAs were analyzed using an Ultimate 3000 Quaternary UHPLC coupled to a Q Exactive Plus mass spectrometer operating in the positive ion mode with a heated electrospray ionization mark II (HESI-II) probe in an Ion Max Source housing. Samples were kept in a temperature-controlled autosampler at 6 °C. Acyl-CoA separation was accomplished by a Waters HSS T3 2.1 × 150 mm column using a three solvent system: (A) 5 mM ammonium acetate in water; (B) 5 mM ammonium acetate in 95/5 ACN/water (v/v); and (C) 80/20/0.1 ACN/water/formic acid (v/v/v), with a constant flow rate of 0.2 ml/min. Gradient performance was as follows: 98 % A and 2 % B for 1.5 min, 80 % A and 20 % B at 5 min, and 100 % B at 12 min; 0.3 ml/min flow at 100 % B at 16 min; and 0.2 ml/min flow at 100 % C at 17 min, held to 21 min, then re-equilibrated at 0.2 ml/min flow at 98 % A and 2 % B from 22 to 28 min. Flow from 4 to 18 min was diverted to the instrument. The mass spectrometer operating conditions were as follows: auxiliary gas 10 arbitrary units (arb), sheath gas 35 arb, sweep gas 2 arb, spray voltage 4.5 kV, capillary temperature 425 °C, S-lens RF level 50, auxiliary gas heater temperature 400 °C, and in-source CID 5 eV. Scan parameters conditions were alternating full scan from 760 to 1800 m/z at 140,000 resolution and data-independent acquisition (DIA) looped three times with all fragment ions multiplexed at a normalized collision energy (NCE) of 20 at a resolution of 280,000. An isolation width of 7 m/z with an offset of 3 m/z was used to capture all relevant isotopologues for targeted acyl-CoA thioesters. Data was processed in Xcalibur 3.0 Quan Browser and TraceFinder 4.1 (Thermo) and statistical analysis was conducted in Excel 2016 (Microsoft) and GraphPad Prism software (v7) (GraphPad, La Jolla, CA). Relative abundance and isotopologue percentage were calculated for each metabolite. Relative abundance of acyl-CoA metabolites was calculated as the sum of the area of all measured isotopologues/the area of the internal standard. For acyl-CoAs where a standard was available, the ratio of the sum of the area of all measured isotopologues/the area of the internal standard was used to generate standard curves which were linear and sample values were interpolated from this curve. Isotopologue enrichment (%) was computed using Fluxfix (http://fluxfix.science), a free web-based calculator (22) using unlabeled samples for experimental isotopic correction.

## Results

### 2M2PE-CoA can be generated via purified protein extracts using propionyl-CoA and reducing equivalents

Previously, we demonstrated that treating HepG2 cells with propionoate creates a dose and time-dependent formation of 2M2PE-CoA (16). To confirm the pathway of 2M2PE-CoA production, HepG2 protein extracts were purified using FPLC separation and incubated for 2 hours with 100 µM propionyl-CoA, 1 mM NADH, and 1 mM NADPH. Of the 156 FPLC fraction collected, samples 90 – 100 converted propionyl-CoA into 2M2PE-CoA as confirmed by the production of both the 2M2PE-CoA product and a putative intermediate, 3-hydroxy-2-methyl-pentanoyl-CoA, identified in the previously proposed mechanism for the conversion of two molecules of propionate into trans-2-methyl-2-pentenoyl-CoA (**Fig. S1-S2**) (16). The results were verified with both negative and positive controls. Negative controls that yielded no product or the intermediate included incubating the protein extract without substrates and incubating the substrates in buffer only. The positive controls which yielded both 2M2PE-CoA and putative 3-hydroxy-2-methyl-pentanoyl-CoA included incubating the entire protein extract, collected prior to FPLC separation, with the substrates, and incubating HepG2 cells in media supplemented with 100 µM propionate for 1 hour (**Fig. 2B**). To confirm the use of NADH/NADPH as a cofactor in the conversion of propionyl-CoA into 2M2PE-CoA, we monitored changes NADH/NADPH concentration by measuring the absorbance at 340 nm. Both NADH and NADPH absorb light at 340 nm, while the oxidized forms (NAD^+^ and NADP^+^) do not (23) and FPLC samples 90-100 exhibited a decrease in NADH/NADPH concentration, correlating with the production of 2M2PE-CoA in these fractions (**Fig. S1**).

**Figure 2:**
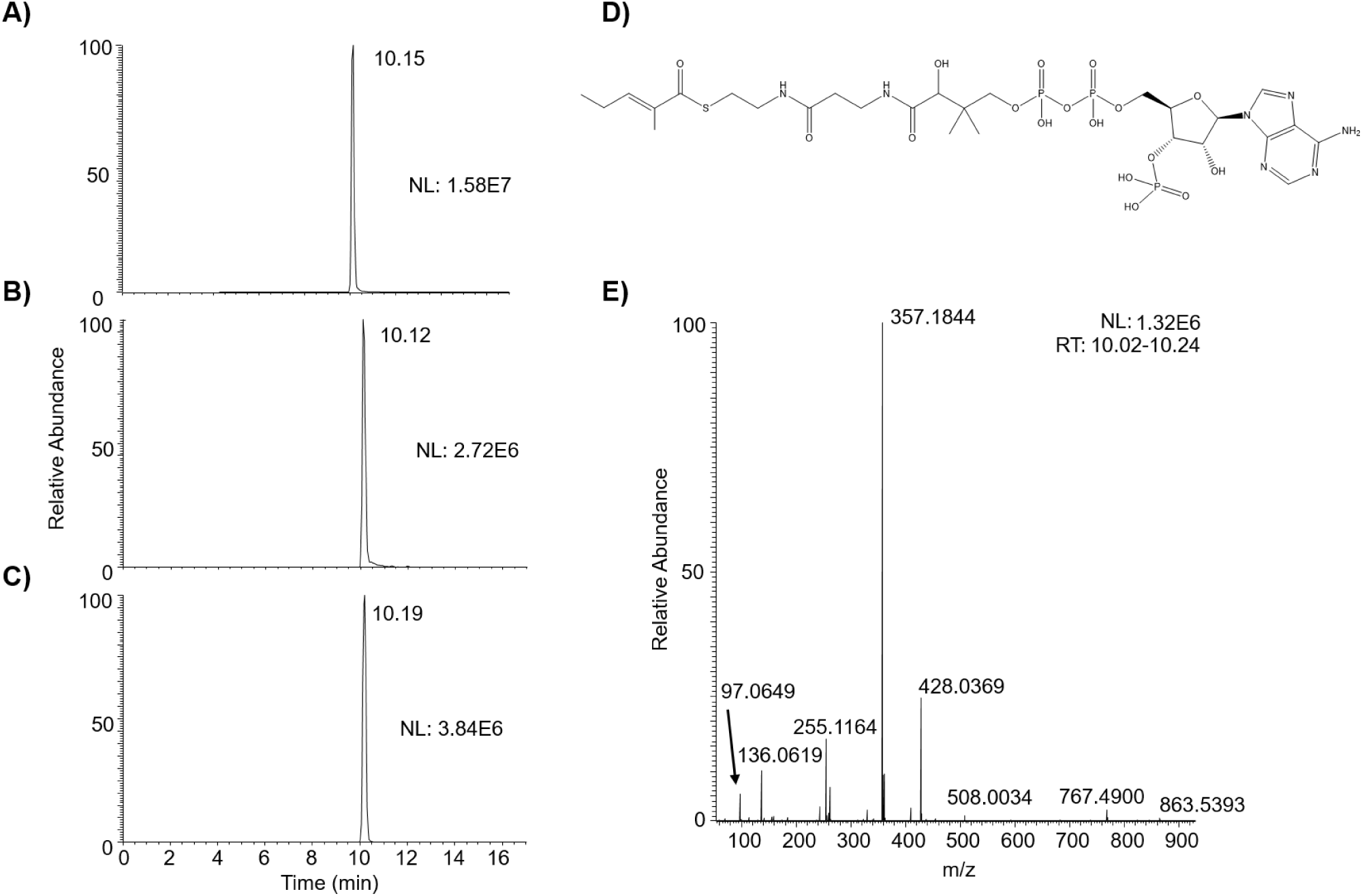
Comparison of synthetic and biologically generated 2M2PE-CoA. Co-elution of A) synthetic standard, B) 2M2PE-CoA produced in HepG2 cells treated with 100 µM propionate, and C) 2M2PE-CoA produced in an active fraction of HepG2 protein extract incubated with 100 µM propionyl-CoA, 1 mM NADH, and 1 mM NADPH. D) Chemical structure of 2M2PE-CoA. E) LC-MS/MS of the synthetic standard.

### Analytical confirmation of the structure of 2M2PE-CoA

To confirm the identification of 2M2PE-CoA generated biologically, we produced a synthetic 2M2PE-CoA standard. LC-MS/HRMS data had matching co-elution and fragmentation of the synthetic standard (**Fig. 2A**) with biologically generated 2M2PE-CoA from both HepG2 cells (**Fig. 2B**) and purified protein extract (**Fig. 2C**). For the HRMS we observed a [MH]^+^ ion at m/z= 864.1811 which matches the theoretical m/z (864.1800) for C_27_H_45_N_7_O_17_P_3_S (Δppm = 1.274) (**Fig. S3**). The MS/MS included the common fragment containing the acyl-group of the [MH-507]^+^ neutral loss distinctive to acyl-CoAs, as well as other fragments reflecting parts of the CoA moiety and the acyl-group reported elsewhere (**Fig. 2E**) (24). Co-elution was further confirmed by HepG2 cells incubated with propionate, 2M2PE, ^13^C_1_ propionate reflecting a predominant +2 mass shift (incorporating 2 units of propionate), or a ^13^C_3_ 15N_1_ label on the pantothenate derived moiety of the CoA backbone reflecting a +4 mass shift (**Fig. S4**). I added

### 2M2PE-CoA forms from valine metabolism via propionyl-CoA in multiple murine tissues

Propionyl-CoA is an intermediate in a large number of metabolic pathways, including branched chain amino acid metabolism. Since our previous work has mostly used propionate directly as a tracer in *ex vivo* settings, we aimed to test the in vivo potential of this pathway while examining the potential of more diverse substrates to contribute carbon to 2M2PE-CoA. To understand the contribution of valine metabolism to potential formation of 2M2PE-CoA, we examined the abundance (pool size) and isotopologue enrichment from infused ^13^C-valine in mice from a previously described study (18). In murine tissues, acetyl-CoA was highly abundant in the liver (mean of 0.77 pmol/mg) and heart (mean of 0.50 pmol/mg), with relatively less abundance on a per mg of tissue basis in muscle (mean of 0.05 pmol/mg) and white adipose tissue (**Fig. 3A and Fig. S5**). Succinyl-CoA demonstrated a similar trend relative to acetyl-CoA across tissues but was less abundant than acetyl-CoA (**Fig. 3B**). Propionyl-CoA was highly variable across tissues, with molar amounts per mg tissue similar to succinyl-CoA (**Fig. 3C**). Unfortunately, insufficient pancreatic tissue remained for analysis in this study.

**Figure 3.**
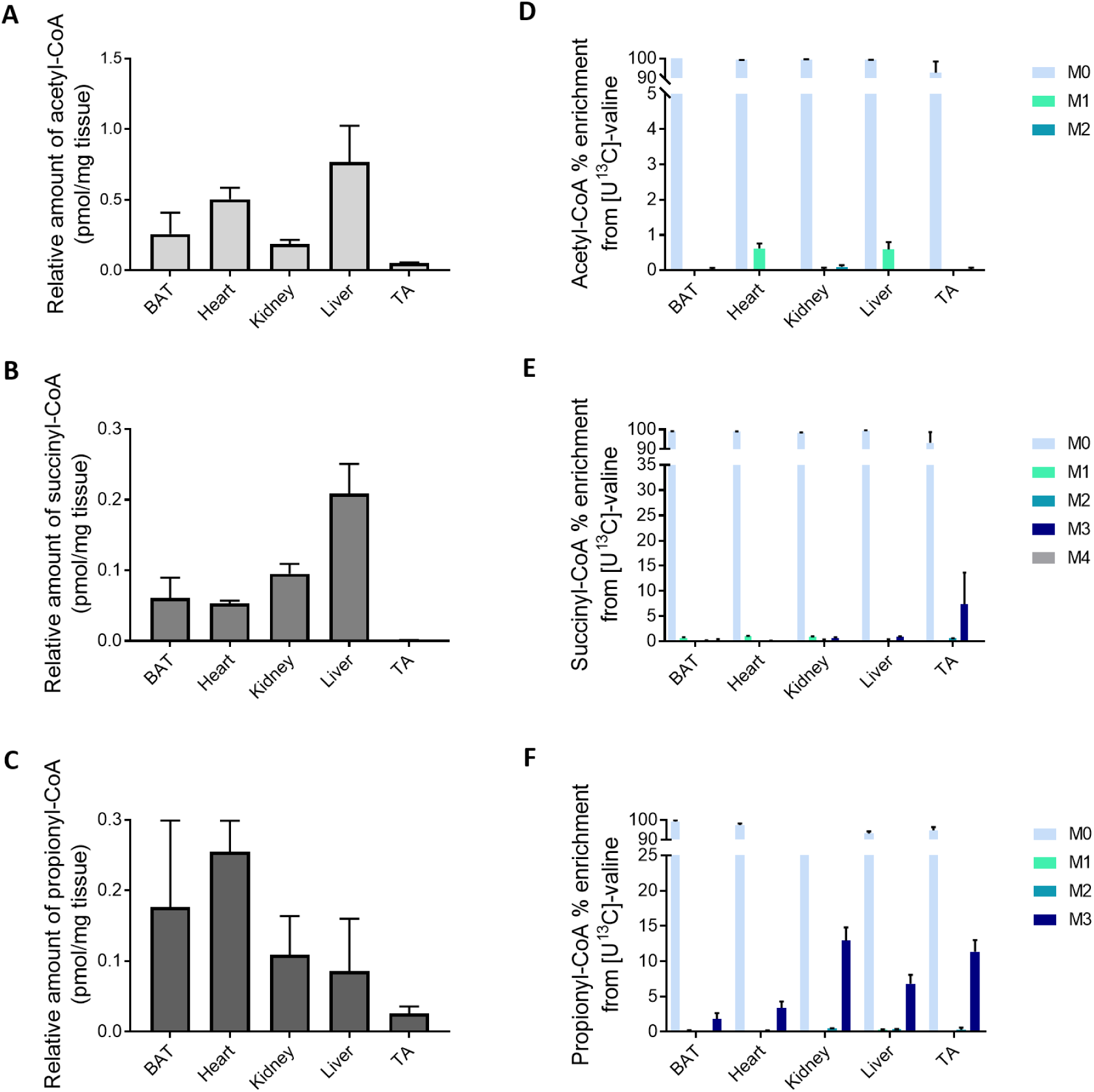
Acyl-CoA abundance and generation from valine varies by tissue type. Relative amounts (pmol/mg tissue wet weight) of (A) Acetyl-CoA, (B) Succinyl-CoA, and (C) Propionyl-CoA, and Isotopologue Enrichment (%) for (D) Acetyl-CoA, (E) Succinyl-CoA, and (F) Propionyl-CoA in BAT, heart, kidney, liver, and TA after 120 minute infusion of ^13^C-valine at a physiologically relevant steady state concentration in conscious mice.

To examine the fate of the infused ^13^C-valine, we determined the isotopologue enrichment of major short chain acyl-CoAs in the analyzed tissues. Isotopologue enrichment in acetyl-CoA revealed that only a relatively small amount of the acetyl-CoA pool in the examined tissue was derived from ^13^C-valine (**Fig. 3D**). This can likely be explained by both the relatively larger pool of acetyl-CoA as well as the other major substrates for acetyl-CoA including glucose and fatty acids. Isotopologue enrichment analysis of succinyl-CoA across tissues suggests some utilization of ^13^C-valine in muscle, with an average of 7.31% in M3 enrichment in TA (**Fig. 3E**). Propionyl-CoA was variably enriched across tissues for the M3 isotopologue, with the kidney being the most highly labeled tissue and containing a mean of 12.93% enrichment (**Fig. 3F**). Since propionyl-CoA is an intermediate of valine catabolism and all three carbons in the acyl-chain of propionyl-CoA can be derived from the carbon backbone of valine in this pathway, the combination of these labeling patterns strongly suggests breakdown of valine in the TCA cycle. These results strongly support the interpretation from Neinast, *et al*. (18) that the muscle is a large oxidizer of branched chain amino acids. This conclusion was robust even though we did not calculate enrichment values based on plasma valine labeling in a manner similar to Neinast since those calculations were already done for TCA cycle intermediates and we are interested primarily in the fate of the valine derived propionyl-CoA. In sum, these results also suggest that valine is catabolized in the kidney to propionyl-CoA but is not (under the conditions studied) oxidized via the anaplerotic sequence into the TCA cycle through succinyl-CoA, which was described broadly as non-oxidative fates previously. Alternatively, this suggests that other substrates provide a sufficiently large proportion of the succinyl-CoA pool that labeling from valine was not observed. We were therefore interested if, especially in the kidney, the M3 enrichment would appear in 2M2PE since it the tracer did not appear as enrichment in a substantial amount of the succinyl-CoA pool.

In previous studies, we had examined the formation of the novel metabolite 2M2PE from propionate *ex vivo* and in cells (16). Since we observed variable amounts of labeling into propionyl-CoA, we used this experiment to investigate the fate of valine derived propionyl-CoA into 2M2PE-CoA. 2M2PE-CoA was variably abundant across tissues (**Fig. 4A**) and displayed a similar trend to the relative abundance of propionyl-CoA (**Fig. 4A vs 3C**). Calculated spearman rank indicated a strong significant correlation between the amounts of propionyl- and 2M2PE-CoA found in C57/blk6 (r= 0.6429, p= .021), db/db (r= 0.7206, p <.001), and both mice groups when analyzed together (r= 0.734, p <.001) (**Fig. 4B-D**).

**Figure 4.**
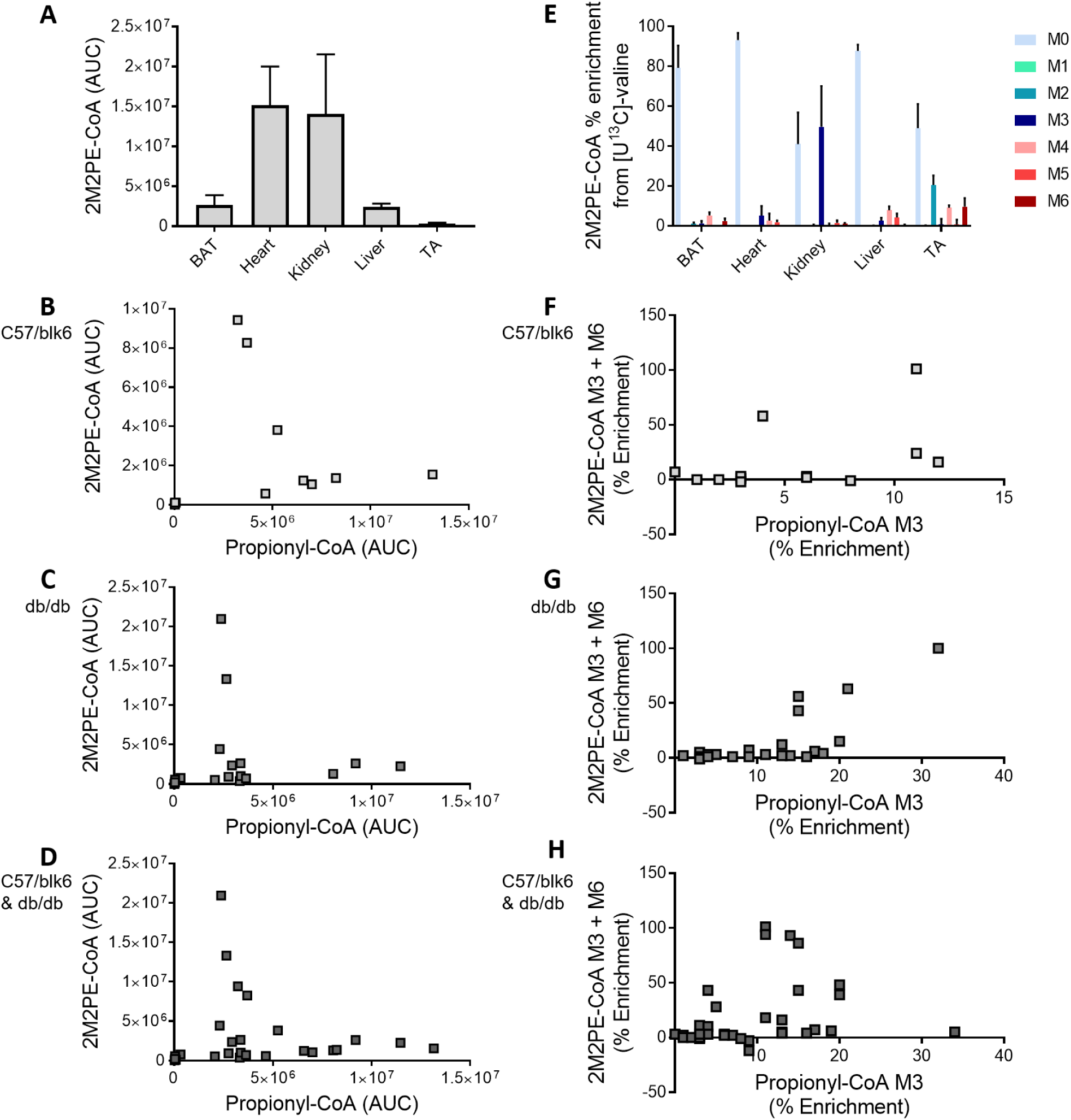
2M2PE-CoA varies in amount and enrichment from valine catabolism across tissues and displays a similar trend to propionyl-CoA in normal and diabetic mice. (A) Area under the curve and (E) Isotopologue enrichment of 2M2PE-CoA labeling incorporating the ^13^C-label from infused ^13^C-valine in murine tissues. Spearman correlation (r) was calculated for the indicated relationships in (B,F) C57/blk6 (n=13), (C,G) db/db (n=22), and (D,H) both mouse groups combined (n=35). Strong significant correlations were found in all variables, with the expectation of (F) 2M2PE-CoA M3 + M6 and propionyl-CoA M3 enrichment % in normal mice (r= 0.4542, p=.120).

Striking isotopologue enrichment of M3 of 2M2PE-CoA in the kidney (mean of 49.67%) suggested half the molar amount of the analyte in that tissue was derived from ^13^C-valine (**Fig. 4E**). We noted that this is higher than the enrichment of propionyl-CoA in the tissue-indicating that there is an as yet unexplained preferential fate for the propionyl-CoA formed from valine catabolism to form 2M2PE-CoA. There was a strong and statistically significant correlation for the association between the enrichment % of propionyl-CoA M3 and 2M2PE-CoA M3+M6 in leptin deficient diabetic db/db mice (r= 0.6355, p= .001) and when data for C57/blk6 control and db/db were combined (r= 0.5443, p <.001) (**Fig. 4G-H**). There was a weaker, not statistically significant association, when calculating this enrichment correlation in C57/blk6 control mice (r= 0.4542, p=.120) (**Fig. 4F**). ^13^C-labeling of 2M2PE-CoA from ^13^C-propionate was also enriched in the heart and to a lesser extent BAT, the liver and muscle (**Fig. 4E**). In BAT, heart, and liver tissues, isotopologue enrichment indicated that 2M2PE was variably but only somewhat derived from valine catabolism. gWAT showed a diffuse enrichment from M0 to M6; however, a major caveat of analysis in this tissue is the considerably lower relative abundance of 2M2PE-CoA in the tissue (**Fig. 4A and Fig S6)**. The sum of this data suggests that 2M2PE generation and propionyl-CoA levels are strongly linked, with further work required to delineate the effects of other metabolic perturbations on this pathway. Additionally, the different patterns across tissues suggest other tissue dependent pathways or compartmentalized metabolic pathways that may influence the preferential substrates and fates of this 3C pathway *in vivo*. The existence of M+3 suggests, and M+6 proves, that the 6-carbon acyl-chain from 2M2PE-CoA can come entirely from propionyl-CoA, recapitulating our previous observations *ex vivo* with propionate with an *in vivo* demonstration of this pathway from valine.

During valine infusions, db/db mice achieved steady stable labeling at nearly twice the amount (40-45%) compared to C57/blk6 control mice (22-27%) to compensate for their larger body size (**Fig. 5A-C**) (18). Db/db mice showed reduced capacity of valine disposal compared to control mice. Increased % levels of valine labeling show an increased utilization of carbons into downstream intermediates for three-carbon metabolism succinyl-CoA M3, propionyl-CoA M3, and 2M2PE-CoA M3+M6. In controls, incorporation of endogenous valine was very low in succinyl-CoA M3 (0-1%); however, diabetic mice showed a slight increase in enrichment at ∼40-45% valine labeling in BAT, heart, kidney, and liver (0-4%) and scattered distribution of gWAT (0-60%) and TA (3-45%) (**Fig. 5A and Fig. S7**). Propionyl-CoA M3 and 2M2PE-CoA M3+M6 displayed a similar trend to succinyl-CoA M3 with more noticeable changes (**Fig. 5A-C**). Formation of propionyl-CoA M3 and 2M2PE-CoA M3+M6 also increased in a dose-dependent manner with increasing valine treatment (**Fig. 5A-C**).

**Figure 5.**
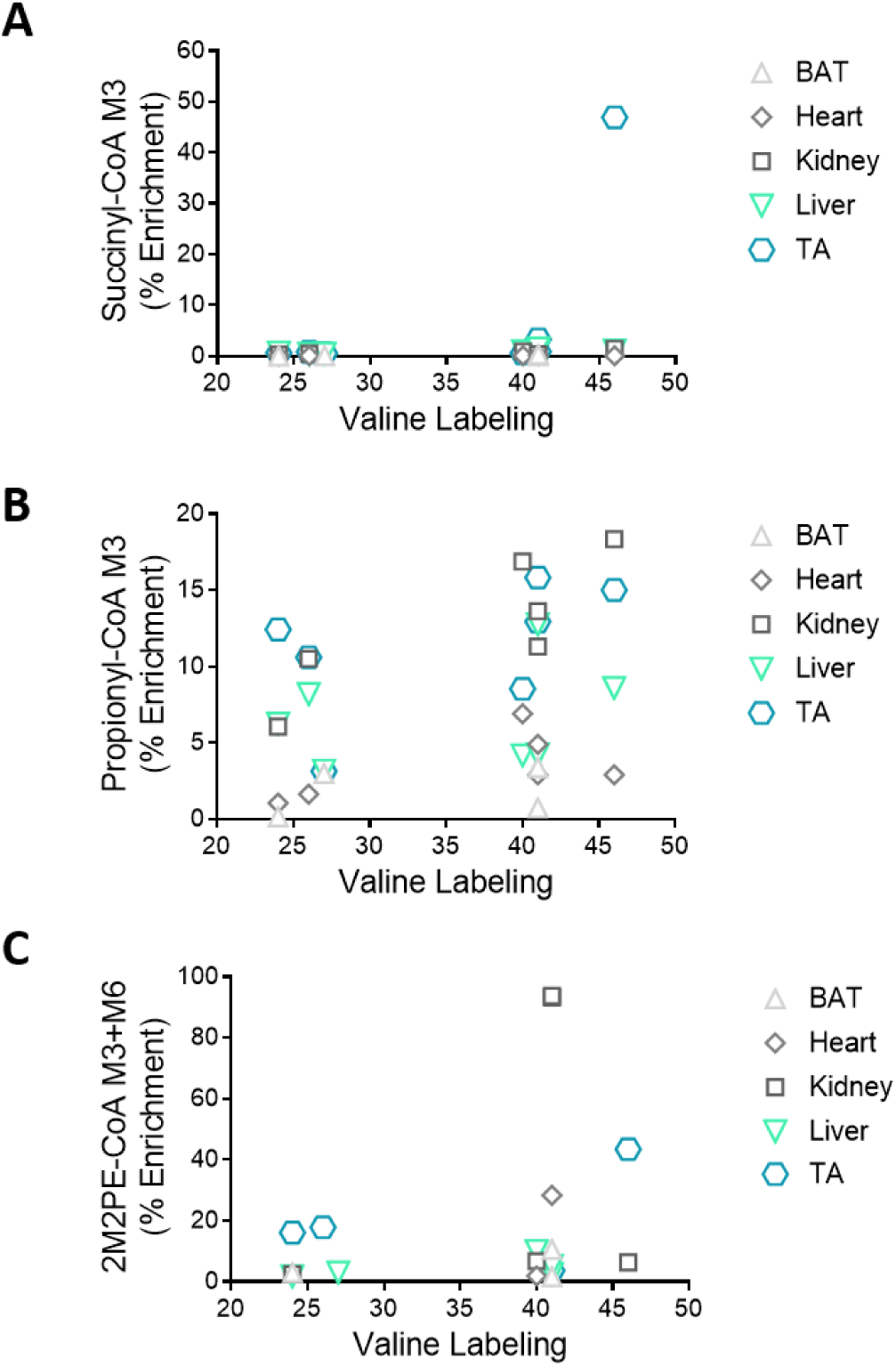
Increased valine labeling correlates with 3C metabolism. Data shows labeling (%) of (A) succinyl-CoA M3, (B) propionyl-CoA M3, and (C) 2M2PE-CoA M3+M6 from plasma valine after constant infusion of [U-^13^C]-valine for 120 min in conscious mice.

### 2M2PE-CoA is generated from propionate in human myocardial tissue

To test if the 3C to 6C pathway extended to human tissue metabolism, we performed a time course study incubating heart tissues in Tyrode’s buffer with 5 mM unlabeled glucose and 100 µM [^13^C_3_]-propionate as a stable isotope tracer. This resulted in apparent pseudo-steady state within 15 minutes, and distinct labeling patterns for acetyl-, succinyl-, propionyl-, and 2M2PE-CoAs (**Fig. 6A-D**). Metabolism via propionyl-CoA provides an anaplerotic pathway forming succinyl-CoA in the TCA cycle (8). If the formation of 2M2PE-CoA was dependent on entry of propionate into the TCA cycle, we might expect to see enrichment in repeating units of M1/M2 (e.g. M4) deriving from acetyl-CoA (C2 metabolism) instead of units of M3/M6 from propionyl-CoA. MID analysis of acetyl-CoA revealed no detectable incorporation of carbon from [^13^C_3_]-propionate, although the relatively large pool size of acetyl-CoA does not preclude the possibility that a small amount of labeling did occur (**Fig. 6A**). Very little M1/M2 succinyl-CoA labeling (both approximately 1-2% enriched) was observed; however, M3 labeling present in succinyl-CoA (around 3-5%) likely arose from anaplerosis into the TCA cycle via propionyl-CoA (**Fig. 6B**). For propionyl-CoA, heart tissues treated with [^13^C_3_]-propionate revealed the anticipated M3 enrichment (**Fig. 6C**). Isotopologue analysis demonstrates that 2M2PE-CoA is not coming from C2 metabolism since labeling in M2+M4 isotopologues is less than M3+M6 suggesting 3C anabolism (**Fig. 2D**). This observation confirms incorporation of propionate into 2M2PE-CoA since M3/M6 labeling is detected throughout apparently healthy human heart *ex vivo* (**Fig. 6B-D**). The generation of the M3 and the M6 isotopologues of 2M2PE-CoA in absence of acetyl-CoA M1 or M2 labeling confirms that the [^13^C_3_]-propionate is not required to first enter the TCA cycle or that the 2M2PE-CoA is derived by canonical fatty acid synthesis via acetyl-CoA.

**Figure 6.**
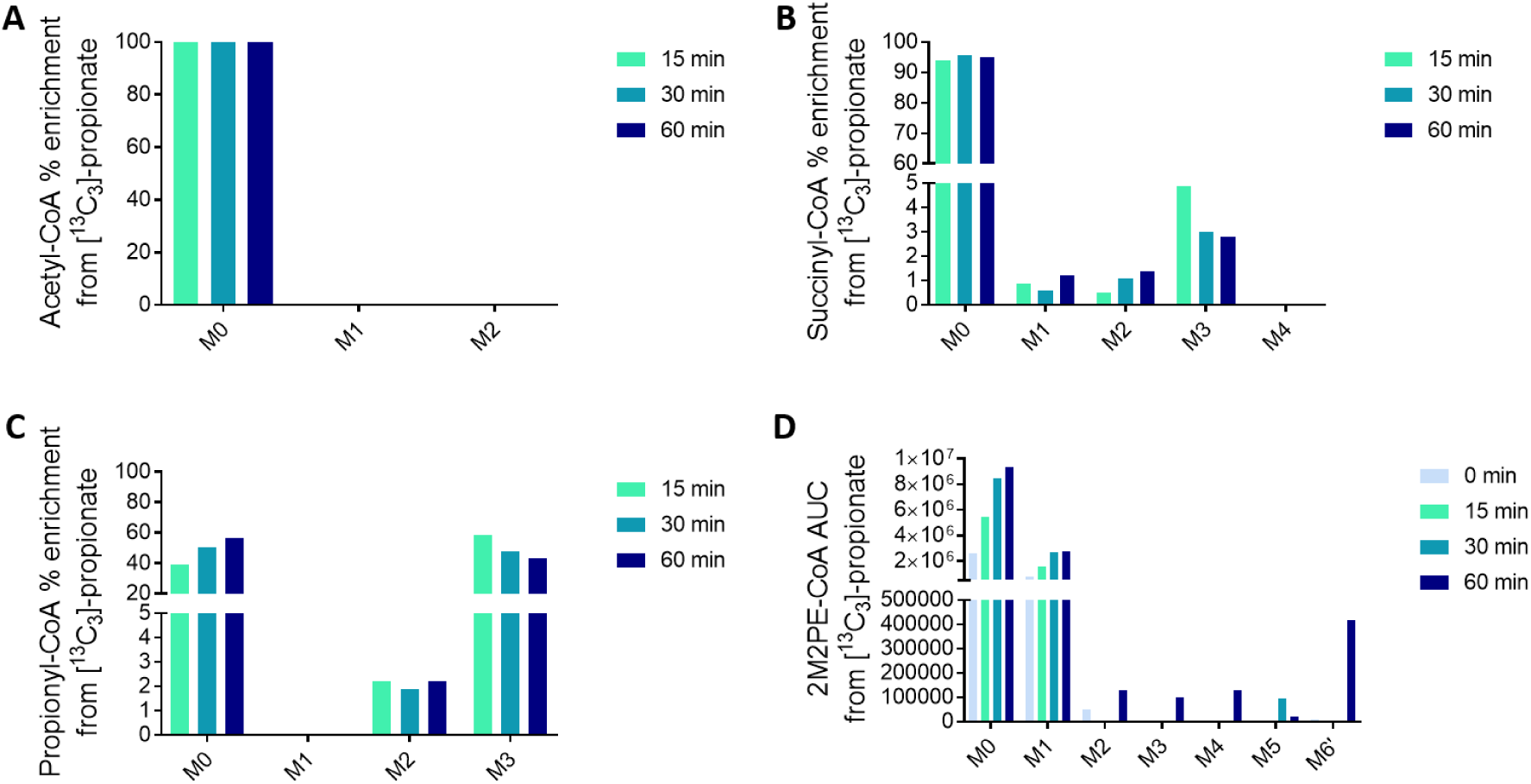
2M2PE-CoA metabolism by propionyl-CoA is not dependent on entrance carbon to the TCA cycle. Time course of generation of (A) Acetyl-CoA, (B) Succinyl-CoA, (C) Propionyl-CoA, and (D) 2M2PE-CoA by LC-MS/HRMS analysis following treatment of heart tissues with 100 µM [^13^C_3_]-sodium propionate in Tyrode’s solution containing 5 mM glucose.

## Discussion

Mammalian anabolic metabolism is traditionally thought to work in units of 1-and 2-carbons. It is believed that mammals lack the capability to utilize three-carbon units of propionate for anabolic reactions, instead shunting excess propionyl-CoA (through D-methylmalonyl-CoA then L-methylmalonyl-CoA) to generate succinyl-CoA as a precursor to other intermediates in the TCA cycle (8, 9). We show here that 3C metabolism occurs in intact human heart from propionate, and in live conscious mice under physiological conditions from valine. We demonstrate that two molecules of propionyl-CoA derived from propionate or valine catabolism, may condense to form a six-carbon 2M2PE-CoA. This replicates our previous findings with both ^13^C-and ^2^H-labeled propionate giving rise to labeling into 2M2PE-CoA that requires a stoichiometry of 2 propionate molecules without metabolism through acetyl-CoA or 1-carbon metabolites in multiple human and murine cell lines, as well as isolated human platelets (16). Importantly, the experiments with fractionated proteins demonstrates that this reaction occurs from propionyl-CoA, NAD(P)H, and the still unknown enzyme(s).

In humans, sources of propionate include the gut microbiota and propionate from food additives. As a food additive, propionate is a used as an anti-microbial and is “generally regarded as safe”. Recently, consumption of this additive has been linked to deleterious metabolic effects. Tirosh *et al*. showed that propionate consumption increased endogenous glucose production leading to hyperinsulinemia via increased norepinephrine, glucagon, and fatty acid-binding protein 4 (FABP4 or aP2) in mice and humans (11). This is a concern, given the addition of propionate in foods, as well as compelling evidence that chronic hyperinsulinemia can drive obesity, the development of type 2 diabetes, and other metabolic abnormalities (25-27). Further elucidation of the 2M2PE-CoA pathway could benefit in unraveling the various metabolic effects of propionate in humans.

Non-oxidative fates of propionate are poorly understood in humans and this poses a challenge in the experimental use of propionate, studies of dietary fates of propionate, as well as the interpretation of pathological propionate metabolism. While oxidative fates of propionate can be quantitatively accounted for by examining tracing into the TCA cycle, non-oxidative fates remain comparatively unexplored. Thus, propionate is commonly used as an isotope tracer to examine TCA cycle metabolism due to tissue and context specific utilization via anapleurosis to succinyl-CoA. Accounting for non-oxidative fates, including but not necessarily limited to 2M2PE-CoA, may improve the interpretation and performance of propionate tracer models.

The propionyl-CoA to *trans*-2-methyl-pentenoyl-CoA metabolic pathway may explain the plethora of branched chain fatty acids and unidentified metabolites that form in the blood and urine of PA patients (15). This route is of particular interest because it generates a relatively non-polar, but more acidic intermediate that can potentially be metabolized to a diversity of branched chain products which may alter the toxicity from high levels of propionic acid (16). Currently, elevated propionylcarnitine (C3) is the primary metabolic finding of PA at diagnosis by newborn screening (28). However, diagnosis by C3 is prone to false-positives, leading to significant diagnostic odyssey until secondary tests can be conducted (29). Furthermore, PA patients can enter metabolic crisis before newborn screening results are available or confirmed, thus earlier biomarkers of PA control are desirable (13). With understanding of *trans*-2-methyl-2-pentenoic acid and clinically observable products, we can identify biomarkers of excess propionic acid metabolism. Future work that identifies the enzymology of this pathway may be warranted to elucidate the functional consequences of metabolism of propionyl-CoA to 2M2PE-CoA.

## Conclusion

Anabolic conversion of 3-carbon propionyl-CoA to 6-carbon 2-methyl-2-pentenoyl-CoA occurs *in vivo* from valine. Conversion of propionate to 2M2PE-CoA was conserved in human hearts. Future work should elucidate the mechanism of control and enzymology responsible for this metabolic pathway.

## Acknowledgements

The authors would like to thank Sanhka “Bobby” Basu for discussions improving the manuscript. This work was funded by NIH grants T32-GM07229 to M.D.N., ADA fellowship 1-18-PDF-144 to S.T., ADA fellowship 1-17-PDF-076 to C.J., Intramural Research Program of NIH (ZIA BC011488-06) NIH grants HL094499 and DK107667 to Z.A., and NIH grants GM132261 and HD092630 to NWS.

