## Supplemental Figures for "Direct anabolic three carbon metabolism via propionate to a six carbon metabolite occurs in human heart and in vivo across mouse tissues"

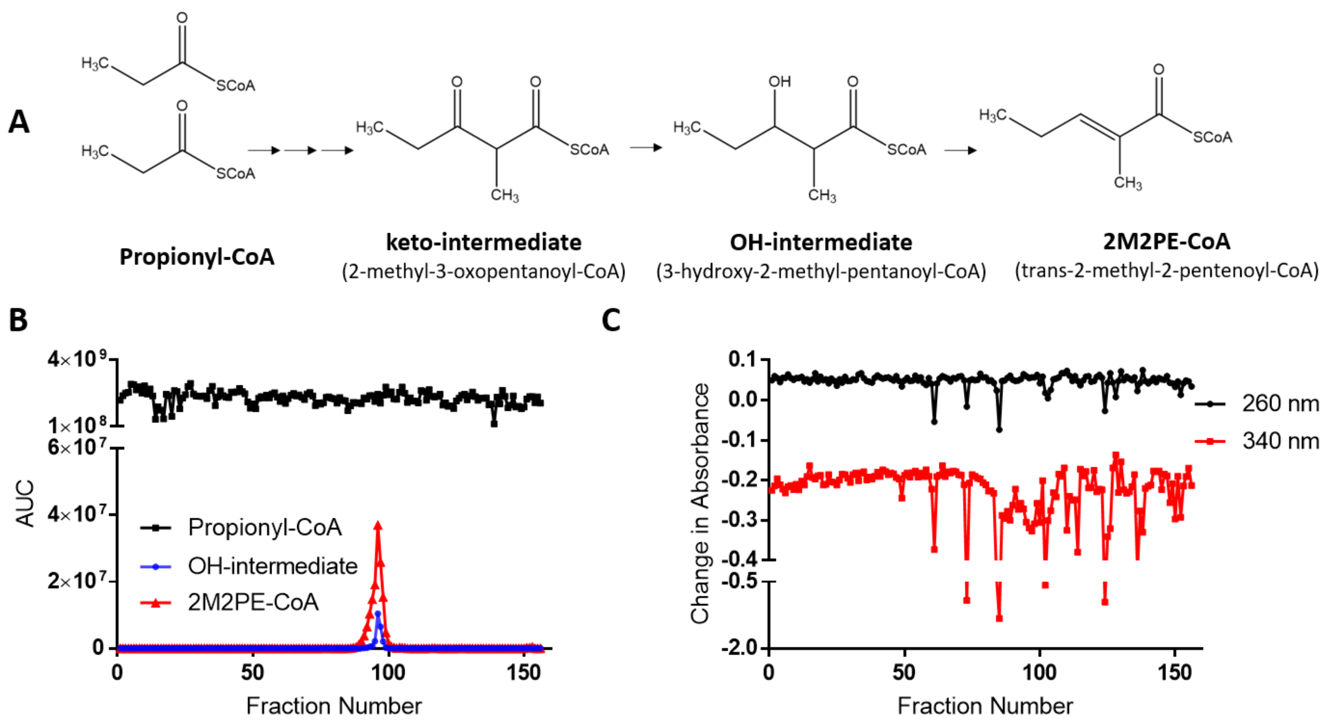


**Figure S1. FPLC purified protein extracts generate 2M2PE-CoA.** A) Abbreviated version of the previously proposed mechanism.[1] B) Production of 2M2PE-CoA (red) and the OH-intermediate (blue) from FPLC protein fractions incubated for 2 hours with propionyl-CoA (black) and NADH/NADPH. C) Change in absorbance following the 2 hour incubation. Both NADH and NADPH absorb light at 340 nm, while the oxidized forms ($NAD^+$ and $NADP^+$) do not.[2] The decrease in absorbance at 340 nm signifies NADH/NADPH oxidation. Absorbance at 260 nm is a negative control because both the oxidized and reduced forms absorb light at this wavelength. The keto-intermediate was not detected due to an interfering peak.


1. Snyder NW, Basu SS, Worth AJ, Mesaros C, Blair IA. Metabolism of propionic acid to a novel acyl-coenzyme A thioester by mammalian cell lines and platelets. J Lipid Res. 2015;56(1):142-50. Epub 2014/11/27. doi: 10.1194/jlr.M055384. PubMed PMID: 25424005; PMCID: PMC4274062.
2. Walker, JRL. Spectrophotometric determination of enzyme activity: alcohol dehydrogenase (ADH). Biochemical Education 1992; 20: 42-43. https://doi.org/10.1016/0307-4412(92)90021-D

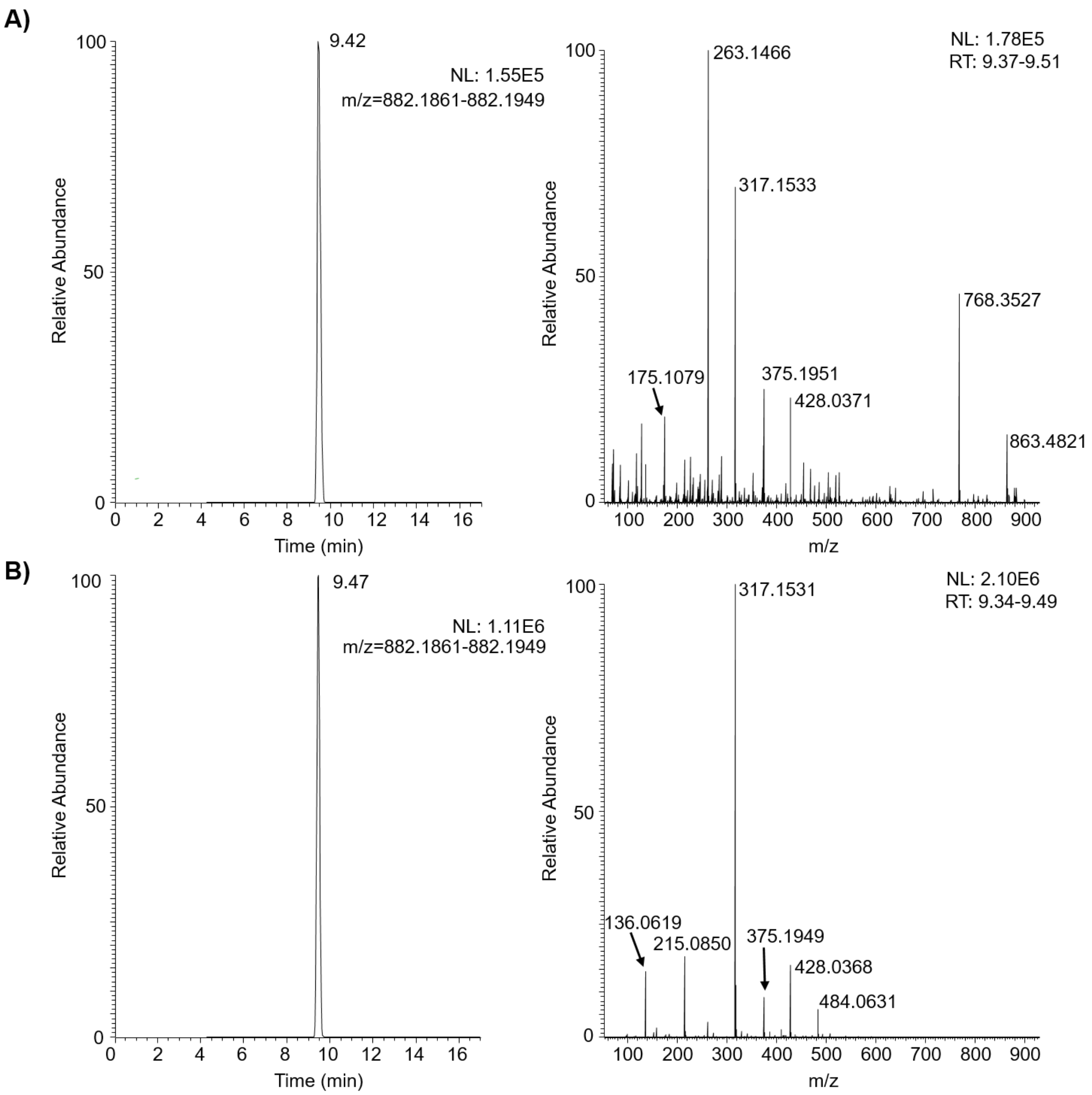


**Figure S2: OH-intermediate (3-hydroxy-2-methyl-pentanoyl-CoA) characterization.** Co-elution and corresponding LC-MS/MS of the OH-intermediate produced in A) HepG2 cells treated with 100 µM propionate for 1 hour and B) an active fraction of the FPLC purified protein extract treated with 100 µM propionyl-CoA, 1 mM NADH, and 1 mM NADPH.

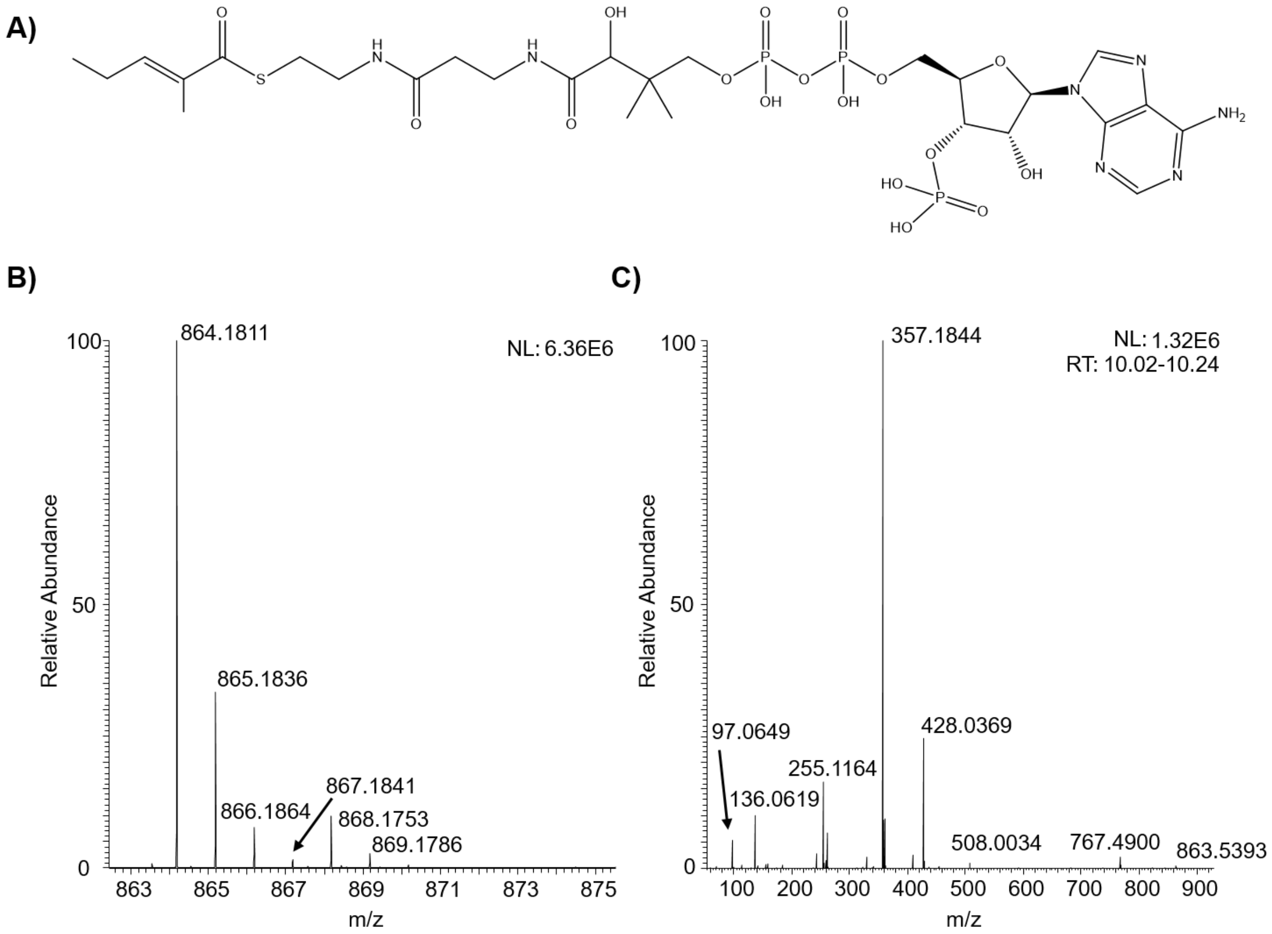


**Figure S3: 2M2PE-CoA synthetic standard characterization**. A) Chemical structure, B) full scan LC-HRMS, and C) LC-MS/MS

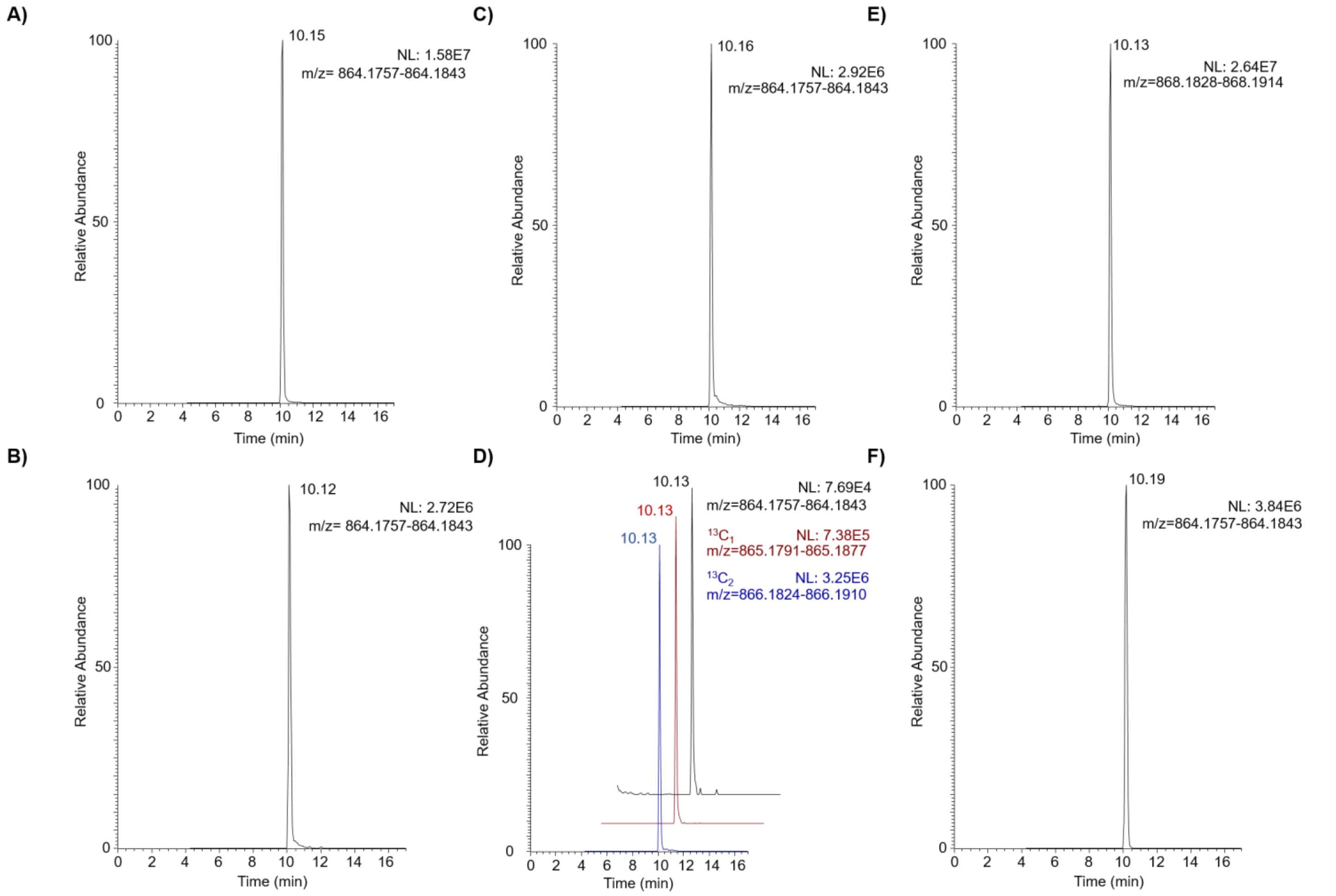


**Figure S4: Co-elution of a synthetic standard and biologically generated 2M2PE-CoA** A) Synthetic 2M2PE-CoA standard. 2M2PE-CoA generated in HepG2 cells treated with B) propionate or C) trans-2-methyl-2-pentenoic acid. D) HepG2 cells treated with $^{13}C_1$-propionoate generated 2M2PE-CoA with a predominant +2 mass shift or a less abundant +1 mass shift from the incorporation of either 2 units or 1 unit of $^{13}C_1$-propionoate. Chromatograms have been offset for clarity. E) SILEC HepG2 cells ([$^{13}C_3{}^{15}N_1$] pantothenate derived labeled)[1] treated with propionate generated 2M2PE-CoA with a +4 mass shift. F) 2M2PE-CoA generated in the active fraction of FPLC purified protein extract treated with propionyl-CoA, and NADH/NADPH.


1. Snyder NW, Tombline G, Worth AJ, Parry RC, Silvers JA, Gillespie KP, Basu SS, Millen J, Goldfarb DS, Blair IA. Production of stable isotope-labeled acyl-coenzyme A thioesters by yeast stable isotope labeling by essential nutrients in cell culture. Anal Biochem. 2015;474:59-65. Epub 2015/01/13. doi: 10.1016/j.ab.2014.12.014. PubMed PMID: 25572876; PMCID: PMC4413507.

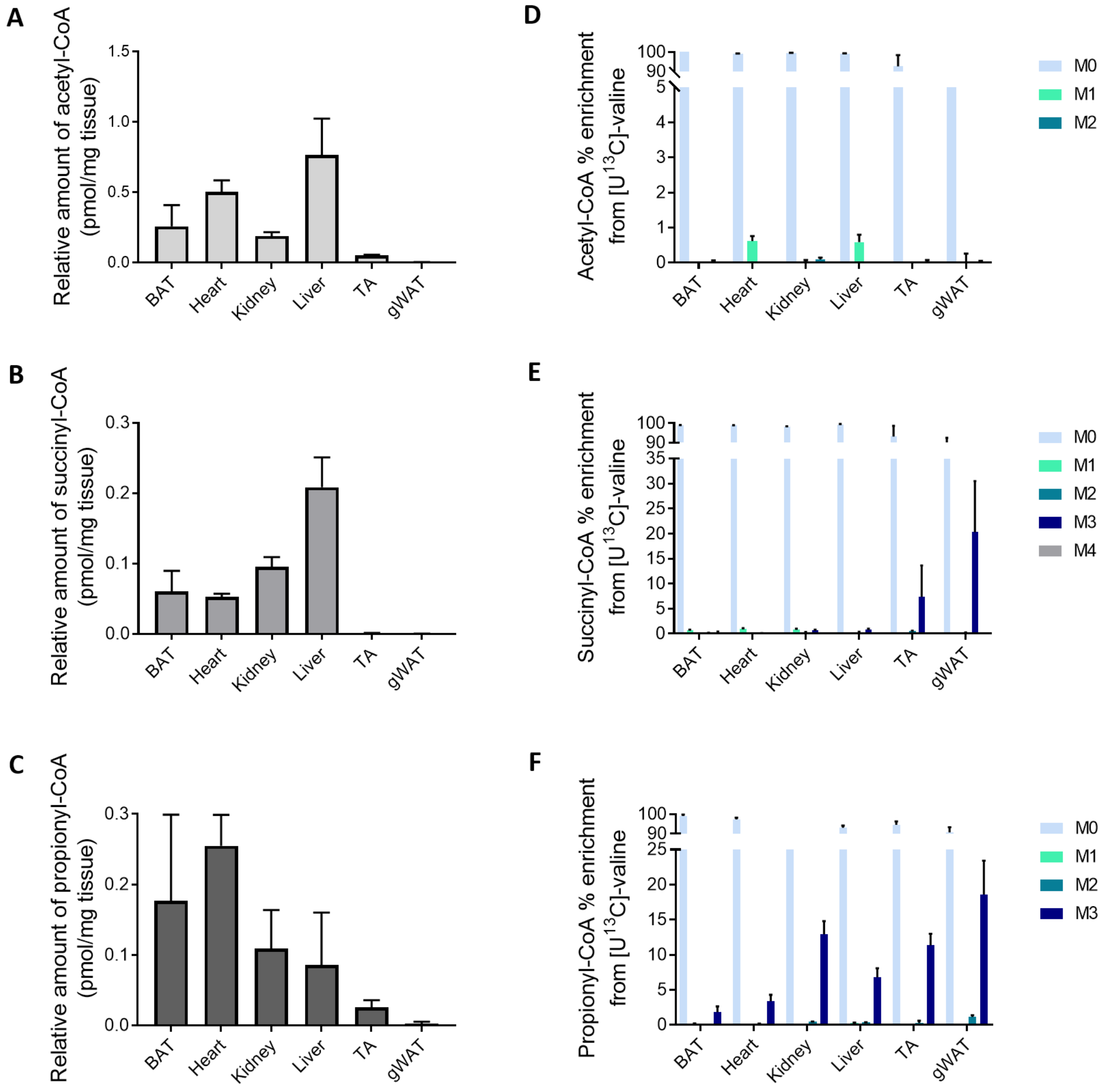


**Figure S5**. **Data from figure 3 with the addition of data from gonadal white adipose tissue (gWAT) tissue.** The low abundance in the gWAT tissue may contribute to a higher contribution of noise to isotopologue analysis, thus interpretation should consider this limitation.

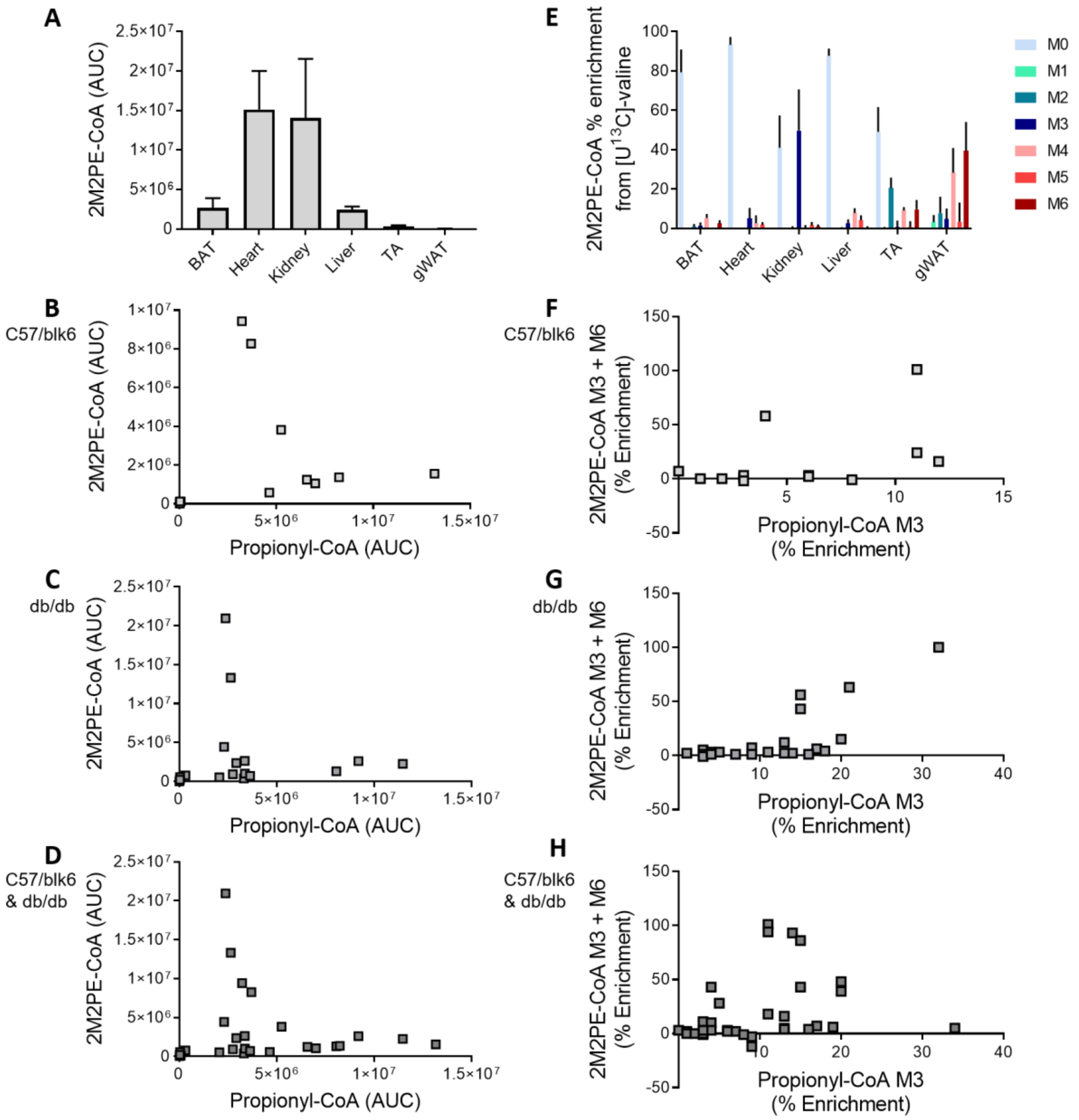


**Figure S6. Data from figure 4 with the addition of data from gonadal white adipose tissue (gWAT) tissue.** The low abundance in the gWAT tissue may contribute to a higher contribution of noise to isotopologue analysis, thus interpretation should consider this limitation.

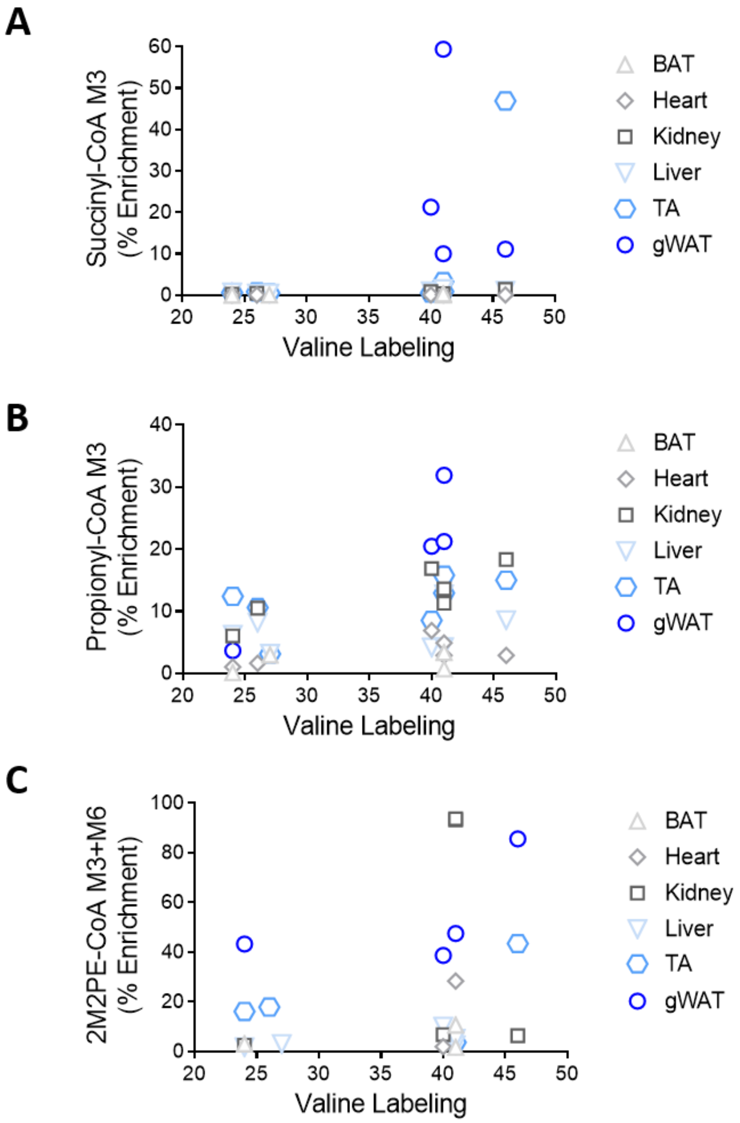


**Figure S7**. **Data from figure 5 with the addition of data from gonadal white adipose tissue (gWAT) tissue.** The low abundance in the gWAT tissue may contribute to a higher contribution of noise to isotopologue analysis, thus interpretation should consider this limitation.
